# PanRes: A database of latent and acquired antimicrobial resistance allowing 3D-based protein homology search

**DOI:** 10.64898/2026.06.22.733705

**Authors:** Martina Vojtková, Martynas Baltušis, Hannah-Marie Martiny, Amulya Baral, Nikiforos Pyrounakis, Attila Beleon, Rebecca Freitag, Anna Pico-Tomàs, Rolf Sommer Kaas, Thomas N. Petersen, Patrick Munk

## Abstract

Antimicrobial resistance databases are central to genomic surveillance, but resistance determinants remain distributed across resources with different scopes, structures, and annotations. We developed PanRes, a curated resistance database of 11,717 genes integrating acquired and latent determinants of antibiotic, biocide, and metal resistance within a unified ontology. We predicted representative protein structures and clustered them by structural similarity, grouping proteins into 505 high-confidence structurally conserved clusters coherent despite sequence divergence. Their structure-guided alignments were used to build Hidden Markov Models (HMMs) for remote homology search. In seven wastewater metagenomes from five European cities, PanRes 3D-based HMMs expanded detection beyond what was possible with BLAST, with 35.3% of retained hits identified only by the HMMs and generally showing greater divergence from known proteins. For beta-lactamases, several proteins retained beta-lactamase-like folds and catalytic geometry despite weak sequence similarity. PanRes is available through an interactive web platform (https://panres.rambio.dk/), a structure-informed resource for exploring the whole resistome.

## Introduction

Antimicrobial resistance (AMR) is a leading cause of global mortality, directly contributing to 1.27 million deaths each year^1^. It has rapidly emerged as a major public health issue in both high- and low-income countries, driven by antibiotic overuse and highlighted by expanding surveillance programmes. Annual AMR-associated deaths are projected to reach 10 million by 2050^2^. Effective surveillance of resistant pathogens therefore depends on the ability to accurately detect and interpret resistance determinants in sequencing data, which in turn requires up-to-date and well-structured AMR reference databases.

Several such resources have been developed, each with its own scope, structure, and criteria for including genes. These differences affect which resistance determinants are represented and how AMR is detected in sequencing data. The Comprehensive Antibiotic Resistance Database^3^ (CARD) offers a highly developed antibiotic resistance ontology (ARO) along with the Resistance Gene Identifier (RGI) tool, which performs AMR detection based on CARD’s reference sequences and Hidden Markov Models (HMMs). ResFinder^4^ exclusively contains clinically relevant AMR genes that have been naturally mobilised (referred to as acquired), and its associated software can perform AMR gene, mutation, and phenotype prediction from sequencing data. ResFinder is complemented by ResFinderFG^5^, which contains AMR genes identified through functional cloning with experimentally confirmed resistance phenotypes that were not yet seen mobilised in nature (referred to as latent genes in this study). MEGARes^6^ uses an acyclic annotation graph structure, which supports statistical analysis and efficient searching. Tools such as AMRFinderPlus^7^ have access to the curated NCBI Reference Gene Database and HMMs to aid in AMR identification. More specialised databases have also been developed, such as BacMet^8^, which focuses on biocide and metal resistance.

However, the use of multiple databases creates challenges, including overlap between entries, inconsistent annotations, and the need to search many resources for the same genes of interest. To reduce this redundancy, we previously integrated genes from several databases, including those described above, into a gene collection of 14,078 unique sequences^9^. Although this collection reduced redundancy at the sequence level, it did not resolve all limitations of existing database-specific resources. Features such as a comprehensive ontology or trained HMMs remained unavailable for many AMR genes, although both are important for the detection and interpretation of resistance determinants. Ontologies standardise relationships between genes, mechanisms, antimicrobial classes, and phenotypes, while HMMs support sensitive detection of more distant homologs.

Since the release of AlphaFold^10^, its structure database has grown to contain over 214 million predicted protein structures as of 2024^11^. Tools such as ColabFold^12^ and Foldseek^13^ have made large-scale protein structure analysis increasingly feasible: ColabFold accelerates structure prediction through faster homology search, while Foldseek enables rapid structural searching and clustering. Together, such tools allow routine use of structural data for the identification of homologs.

Structurally homologous proteins can have large sequence variation due to structural importance and conservation of fold^14^. Because structure is more conserved than sequence, using 3D structures has been suggested as a more reliable option for protein similarity studies^14,15^. This has also been shown for AMR discovery: Ruppé et al. used comparative modelling to identify distant resistance determinants in the human gut microbiome, many with low amino acid identity to known resistance proteins^16^ and Coche-Miranda et al. used structural clustering to distinguish beta-lactamase candidates from structurally similar proteins with other functions^17^.

The use of 3D structural data for HMM construction was explored as early as the mid-to-late 2000s through the encoding of protein backbones with a structural alphabet^18^. These approaches extended HMM methods originally developed for primary sequence data and enabled remote homology detection. More recently, a structural alphabet describing tertiary interactions has shown to be more accurate and information dense^13^, and has also been used to construct HMMs and look for deeper evolutionary relationships^19^.

Here we present PanRes, an antimicrobial resistance database with comprehensive ontology, predicted protein structures, and HMMs based on structural alignment (Figure 1). We demonstrate the use of the HMMs on large metagenomic datasets, highlighting how the resource can be applied to screen and discover putative distant AMR genes and their non-AMR relatives in complex microbiome datasets. PanRes is available through a web interface (https://panres.rambio.dk/) that allows users to browse, filter, visualize, and download database content while exploring interlinked data across the pan-resistome.

**Figure 1.**
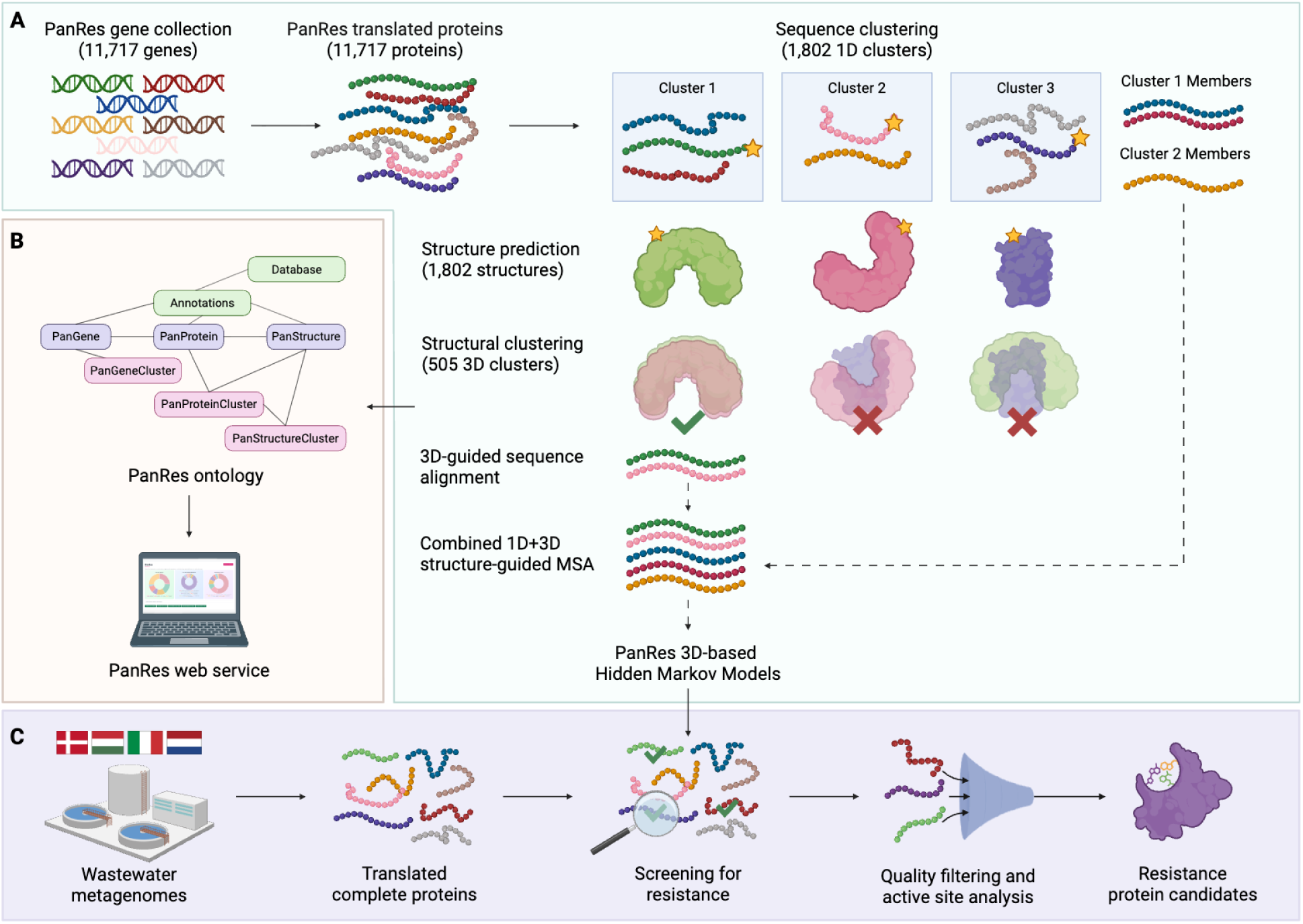
Graphical abstract highlighting the main steps carried out in the study. **A.** Curated resistance gene sequences were translated, clustered at the protein level, and represented by predicted protein structures. Structurally coherent clusters were then used to build alignments and HMM profiles for downstream AMR detection. **B.** Results were organized into a unified ontology and made accessible through the PanRes web service. **C.** The HMMs were applied to wastewater metagenomes to screen for resistance protein candidates, which were further prioritized by catalytic-residue conservation. Created with BioRender.com

## Methods

### Curating the database

The new PanRes database was developed based on the gene collection distributed with ARGprofiler^9^. It consisted of 14,078 unique DNA sequences from eight databases provided as a FASTA file and accompanying metadata table.

Protein-coding sequences were predicted and translated using Prodigal^20^. Because PanRes contains genes from diverse sources, including entries from metagenomic and functional cloning studies, Prodigal was run in both *meta* (anonymous) and *single* (normal) modes.

We refined the results from both modes by (i) excluding sequences with multiple translations, (ii) filtering out sequences without valid start or stop codons, (iii) removing sequences that produced proteins of unexpected length based on an expected nucleotide-to-amino-acid ratio of 3:1, and (iv) merging the remaining sequences from meta and single modes. Using the refined Prodigal output as reference, the original PanRes nucleotide sequences were filtered to retain only entries producing a single complete coding sequence.

### Sequence clustering

The nucleotide sequences were clustered with USEARCH^21^ (v12.0 [*b1d935b*]) using the *cluster_fast* algorithm at 90% sequence identity and 90% coverage to 4,199 gene clusters.

The protein sequences were clustered with CD-HIT^22^ (v4.8.1) with the parameters set at 50% identity and 90% coverage (from here referred to as 1D clusters), as proteins within this range are generally expected to retain the same overall fold^23–25^. Each cluster was represented by a single centroid sequence, the longest sequence in each cluster.

### Protein structure predictions

We predicted 3D protein structures for the centroid sequence of each 1D cluster using LocalColabFold (v1.5.5)^12^. Multiple sequence alignments (MSAs) were generated with *colabfold_search* using environmental sequences and structural templates, and structure prediction was performed with *colabfold_batch* on NVIDIA Tesla V100 16GB GPUs. The highest-ranking predicted structure for each protein was retained for downstream analyses. Database used and prediction parameters are described in Supplementary Methods.

### Structure-based clustering and pairwise structural comparison

The representative structures were clustered based on three-dimensional similarity using Foldseek easy-cluster^13^ (release 10-941cd33), run in global alignment mode with TMalign and clustered on the alignment Template Modeling (TM) score, requiring a minimum alignment coverage of both the query and the target structures.

Several combinations of query/target coverage (75% - 90%) and alignment TM-score thresholds (0.6 - 0.7) were tested (Supplementary table 1). The TM-score threshold of 0.6 was selected as the lower bound, as indicative of shared overall protein fold and structural homology^26^. Based on cluster coherence, singleton-cluster frequency, and intra-cluster structural similarity, we selected 80% coverage and an alignment TM-score of 0.6 for downstream analyses.

Prediction confidence was assessed using per-residue pLDDT values from the retained ColabFold models. Structure predictions with a mean pLDDT <70 were classified as low-confidence^27^, and structural clusters containing only these models were removed from the final structural resource. We further assessed cluster coherence by per-cluster Pfam-A^28^ protein-family consistency (Supplementary Methods).

All-vs-all pairwise comparison was also performed using Foldseek’s easy-search with exhaustive search enabled, mirroring the global TMalign clustering settings. Full Foldseek parameters are listed in Supplementary Methods.

### Clustering on primary and tertiary protein structure to inform Multiple Sequence Alignments

We combined information from both sequence-based (CD-HIT) and structure-based clustering (Foldseek) to generate broader MSAs for each protein family. First, an alignment was generated for each structural cluster using T-COFFEE^29^ (v13.46). T-COFFEE was run in structure-aware alignment mode (Expresso^30^) using the *TMalign_pair* method, which incorporates pairwise structural superpositions generated by TM-align to guide the sequence alignment.

Since the structural clusters were based only on the centroids of 1D clusters, the resulting MSAs were expanded back to include all proteins from the corresponding cluster. The sequences were added to the existing 3D alignments using MAFFT^31^ in *add* mode, ensuring the structure-guided MSA backbone remained unchanged. The final MSA for each protein family was then used to build Hidden Markov Models.

### PanRes Hidden Markov Models

HMM profiles were built from the combined protein MSAs using the *hmmbuild* function from HMMER^32^ (v3.4) with default settings. HMMs of all individual clusters were then merged and the database was prepared with *hmmpress*.

We evaluated the performance of the PanRes HMMs by measuring how accurately each HMM recognized proteins belonging to the structural cluster from which it was built. The full PanRes protein set was searched against the HMM database using *hmmscan*, and predicted assignments were compared with the known structural-cluster membership. For each clustering parameter combination tested, we recorded proteins without an HMM hit as false negatives, proteins assigned to the wrong cluster as false positives, and recall as the proportion of proteins for which the correct HMM appeared as the top- or second-ranked hit (Supplementary table 2).

To reduce the influence of uncertain structural models on HMM construction, HMM profiles were excluded if low-confidence structures comprised more than 10% of their structural representatives.

### An ontology for PanRes

To ensure consistent integration of information from the source databases, we developed the PanRes ontology to structure annotations for AMR genes and all their related information. Following the syntax and schematic conventions used in the ResFinder database^4^, we harmonised the data structures and terminology in the other source databases. The ontology was built in Python (v3.11.2) using owlready2^33^ (v0.44) and metadata translation was performed using pandas^34^ (v1.5.3).

First, the PanRes^9^ metadata table was parsed to extract gene identifiers and source database information. Genes removed during database curation were assigned to the *DiscardedPanGene* class. These were then excluded from further annotation steps, however, remain part of the ontology to allow for comparison with past database versions.

To represent genes, proteins, structures, and their relationships, we defined core ontology classes for *PanGene*, *PanProtein*, *PanStructure*, and their corresponding cluster classes: *PanGeneCluster, PanProteinCluster*, and *PanStructureCluster* (Supplementary table 3). Gene and protein cluster membership was extracted from the USEARCH and CD-HIT output files, respectively. Object properties such as *member_of, translates_to,* and *folds_to* were assigned to define relationships between ontology individuals (Supplementary table 4). Annotation properties were added from the available metadata, including original gene names, accessions, sequence length for individual genes and proteins, and member counts for clusters (Supplementary table 5).

### *E. coli* core gene homology identification

To identify resistance proteins that may represent housekeeping gene homologs, we compared all PanRes protein sequences against the *E. coli* soft-core genome published as part of an *E. coli* pangenome analysis based on 1,324 genomes^35^. The soft-core genome consisted of 3,058 conserved protein families (193,050 sequences) that were present in >95% of analysed genomes. The soft-core and PanRes proteins were then compared using CD-HIT-2D (v4.8.1). Homology thresholds were set to 60% sequence identity, 60% alignment coverage, as these were the parameters identified as optimal by the study. The PanRes proteins that were identified as homologs, meaning clustered together with soft-core genes, were additionally annotated as *is_ecoli_homolog* in the ontology.

### Screening for resistance with PanRes HMMs and BLASTp

To compare structure-based and sequence-based AMR detection, we applied HMMER searches using PanRes 3D-based profiles and BLASTp searches to proteins from seven metagenomic co-assemblies from wastewater treatment plants in Bologna, Budapest, Rome, Rotterdam, and three sites in Copenhagen^36^, being Avedøre (RA), Damhusåen (RD) and Lynetten (RL). Assembly process, binning, and taxonomic classification were described by Becsei & Fuschi et al.^36^. Coding regions were predicted with Prodigal^20^ (v2.6.3), and only complete protein sequences were retained for downstream analysis.

The proteins were searched against the indexed PanRes HMM database without applying an e-value cutoff. The protein database used for BLAST search was built from PanRes protein sequences. Predicted proteins from the metagenome were searched against the database using BLASTp (v2.17.0+) with the following parameters: e-value threshold of 1, low-complexity filtering enabled, and maximum one high-scoring segment pair (HSP) per target. An e-value threshold of 1 was selected to retain weak sequence matches for comparison with HMM-based detections, while keeping the search computationally feasible.

### Comparison of HMM and BLASTp detection sensitivity

Hits with e-values below 1e−5 were classified as high-confidence, and a single best scoring match per query was retained for both methods. Sequences homologous to the *E. coli* soft-core proteome (*is_ecoli_homolog*) were extracted and matched to PanRes structural clusters. BLASTp hits matching these sequences and HMM hits matching structural clusters in which these homologs represented at least 50% of members were removed from downstream analyses.

We compared the results of both methods at the query level. First, we recorded whether each query protein was detected by both methods or by only one method. For HMMER, matches were assigned at the cluster level, while BLASTp targets were considered equivalent if they belonged to the same structural cluster. For queries detected by both methods, we compared the top matching targets assigned by HMMER and BLASTp. We divided the results into four categories: (i) Agreement, where both methods selected the same target; (ii) Weak disagreement, where the top targets differed but both methods still detected the other method’s top target; (iii) Strong disagreement, where only one method detected the other method’s top target; and (iv) Complete disagreement, where the top targets differed and neither method detected the other method’s top target.

To further compare method sensitivity, we used the permissive BLASTp results (e-value < 100) to place the HMM-detected proteins in sequence similarity space. For each protein, we plotted BLASTp percent identity against alignment length and compared the result with the Rost curve^25^, which estimates the sequence identity threshold for structural homology recognition. We also validated the HMM-unique detections by comparing their predicted 3Di representations, which encode the protein structure in structural alphabet, against the high-confidence PanRes structures using Foldseek^13^ (Supplementary Methods).

### Catalytic residue mapping and scoring

Catalytic residues were mapped from the Mechanism and Catalytic Site Atlas^37^ (M-CSA) to PanRes resistance proteins and their corresponding HMMs. Because M-CSA residue positions are defined in Protein Data Bank (PDB) coordinates, SIFTS^38^ residue-level mappings were used to convert catalytic residue positions from PDB to UniProt/PanRes coordinates. We located the corresponding active site positions in the MSAs used for HMM construction and mapped to HMM model columns. Metagenomic hits were scored based on whether the expected catalytic residues were conserved at these columns. In addition to exact amino acid matches, we also calculated a more permissive chemistry-based score by allowing substitutions between residues with conserved functional groups: D↔E (acids), K↔R↔H (bases), N↔Q (amides), and S↔T↔Y (hydroxyl groups).

### Beta-lactamase candidate analysis

Using the PanRes ontology, we identified the beta-lactamase protein structural clusters. These were defined as clusters containing at least 80% of proteins with antibiotic resistance class annotated as *Beta_lactam*. The member proteins were classified to individual Ambler classes^39^ using the Beta-Lactamase Database^40^ (BLDB) to profile the structural clusters. Four large clusters were selected for active-site analysis: PANCL12631_struct (majority Ambler classes B1/B2), PANCL5265_struct (B3), PANCL9032_struct (A/C), and PANCL8238_struct (A/D). Putative beta-lactamase proteins with completely or chemically conserved catalytic sites were analysed further.

To identify plasmid-associated hits, contigs carrying high-confidence beta-lactamase sequences were screened with geNomad^41^ (v1.12.0). Contigs with a plasmid score above 0.7 were annotated as plasmid-associated. Contigs not associated with any mobile genetic element were classified as chromosomal. MAG assignments were taken from the original study^36^. We retained contigs from metagenome-assembled genomes (MAGs) if they were at least 5 kbp in length, while discarding contigs from genome bins not satisfying medium-quality MAG criteria proposed by Bowers et al.^42^ (MIMAG criteria). Unbinned contigs were only included if they exceeded 10 kbp and were plasmid-associated.

Protein sequences were grouped by HMM cluster, aligned against the corresponding HMM profiles, and used to infer phylogenetic trees with IQ-TREE^43^ (v3.1.1) using ModelFinderPlus^44^. The phylogenies were visualized and annotated in iTOL^45^ (Supplementary Methods).

We predicted 3D structures of selected candidates with ColabFold^12^ (v1.6.1) using the same settings as for the PanRes database structures. Active-site visualizations were generated in PyMOL^46^ (v3.1.6.1).

## Results

PanRes integrates curated latent and acquired resistance genes with protein sequences, predicted structures, structural clusters, HMM profiles, and ontology-based annotations. The database is accessible through a web interface where users can browse the ontology and explore linked gene, protein, and structure information (https://panres.rambio.dk/).

### PanRes curation and the structural resistome

We used the original PanRes component of the ARGprofiler pipeline^9^ as the starting point for building our database. 2,361 sequences missing start or stop codon or containing additional flanking sequences were filtered out. These flanking regions were particularly common in entries derived from functional cloning studies. The curated set contained 11,717 gene sequences, each translated into a corresponding protein sequence. The vast majority (11,486) were identically predicted by both Prodigal’s modes, while 41 and 190 were unique to the *single* and *meta* modes, respectively.

The proteins were clustered into high-homology groups (50% identity, 90% coverage) expected to share the same fold, yielding 1,802 distinct clusters, each with a representative centroid sequence. Predicted tertiary structures for these centroids added a structural layer to PanRes and enabled fold-based clustering of the protein families.

Across the tested coverage and alignment TM-score thresholds, 80% coverage and an alignment TM-score of 0.6 gave the best balance between cluster coherence and singleton frequency. These parameters produced 598 structural clusters of variable sizes (mean ∼ 3, max = 189). 75.75% of all 3D clusters contained only a single structure representative (Supplementary table 1). Following confidence filtering, 99 low-confidence structures (pLDDT<70) were excluded. This resulted in the removal of 93 singleton clusters, leaving 505 structural clusters in the final resource.

The resulting structural clusters were highly coherent. Across clusters, the mean intra-cluster alignment TM-score was 0.844 and the mean Local Distance Difference Test (LDDT) was 0.732 (Figure 2A). Even the least coherent cluster had a mean TM-score of 0.628, remaining above the clustering threshold. At the pairwise level, 99.95% of intra-cluster comparisons exceeded the 0.6 TM-score threshold (Figure 2B). In addition, the majority of clusters contained a single Pfam family (87%).

**Figure 2.**
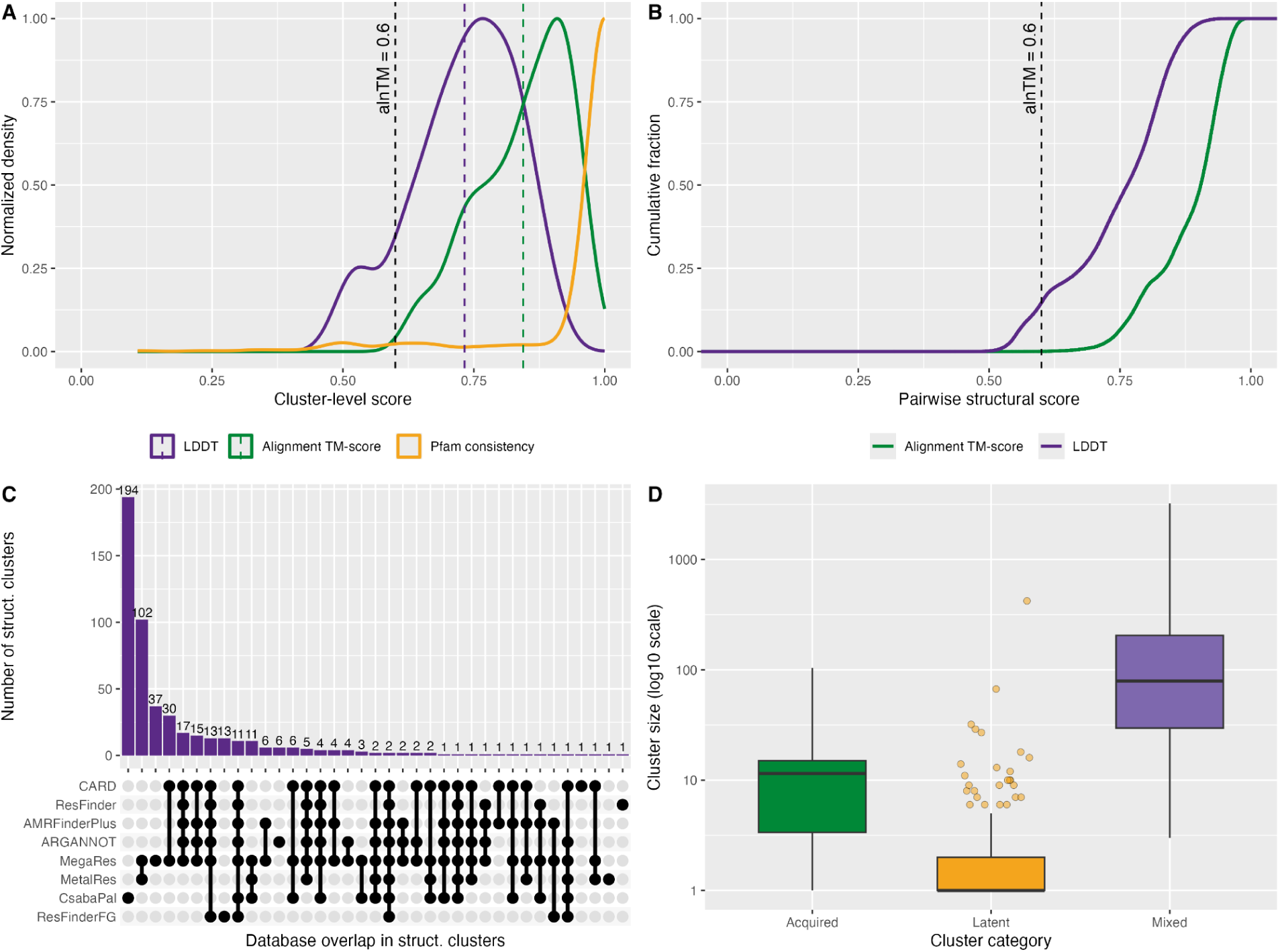
Structural quality and database composition of structural clusters. **A.** Distributions of mean intra-cluster alignment TM-score, mean LDDT, and Pfam consistency (the fraction of proteins per cluster sharing the dominant Pfam annotation) across clusters. Dashed coloured lines mark per-metric averages and the black dashed line the clustering threshold (alnTM = 0.6). **B.** Cumulative distribution of pairwise alnTM and LDDT values within clusters. Most pairwise scores exceed the 0.6 threshold, indicating that clusters remain structurally consistent at the individual pair level. **C.** The database combinations represented within the structural clusters. **D.** Cluster size distribution by resistance origin: acquired (ResFinder support), latent (functional database), or mixed. The box shows the interquartile range (IQR) with the median marked by a horizontal line. Whiskers extend to the most extreme values within 1.5 × IQR, and outliers are shown as jittered points.

Structural clustering based on 3D similarity has the potential to extend protein families beyond conventional sequence-based groups. Foldseek clustering of the representative protein structures revealed that a large number of latent resistance clusters remained structurally distinct: 194 clusters exclusively contained sequences from the functional antimicrobial selection work by Daruka et al.^47^ (hereafter CsabaPal, the corresponding label in PanRes) and 13 clusters exclusively included ResFinderFG sequences, even after expanding the clusters to the full dataset (Figure 2C). This shows that many functionally identified resistance determinants do not share structural similarity with previously characterized AMR proteins.

At the same time, some clusters linked latent and acquired AMR genes. ResFinder genes co-occurred structurally with CsabaPal genes in 19 clusters and with ResFinderFG genes in 26 clusters, suggesting that some clinically known resistance genes are related to broader environmental AMR reservoirs. These links may help place acquired AMR genes in a wider evolutionary context and highlight latent genes that could become clinically relevant. Cluster sizes also differed by origin: latent-only clusters were mostly singletons or small groups (median = 1.0), showing how undersampled especially the latent resistome is. Clusters containing acquired resistance were larger (median = 11.0), while mixed clusters were largest overall (median = 79.5) (Figure 2D).

Most structural clusters contained proteins associated with a single resistance type: antibiotic, biocide, or metal resistance. However, a smaller set of 48 clusters contained proteins annotated with more than one resistance type, dominated by broad-substrate efflux pumps (Supplementary figure 1). This highlights how PanRes can reveal shared resistance potential that is otherwise split across database annotation categories.

### The PanRes ontology

We developed a structured PanRes ontology (Figure 3) that organises the resource’s entities - genes, proteins, predicted structures, clusters, and resistance terms (Supplementary table 3) along with the relationships connecting them (Supplementary table 4). Each entry retains the original name and accession number, so it can be traced back to its original source (Supplementary table 5). Among antibiotic resistance genes, 99.64% were annotated with resistance class, 76.92% with a phenotype, and 44.23% with resistance mechanism. All biocide resistance genes had a resistance class assigned, and all metal resistance genes had a metal phenotype assigned (Supplementary table 6).

**Figure 3.**
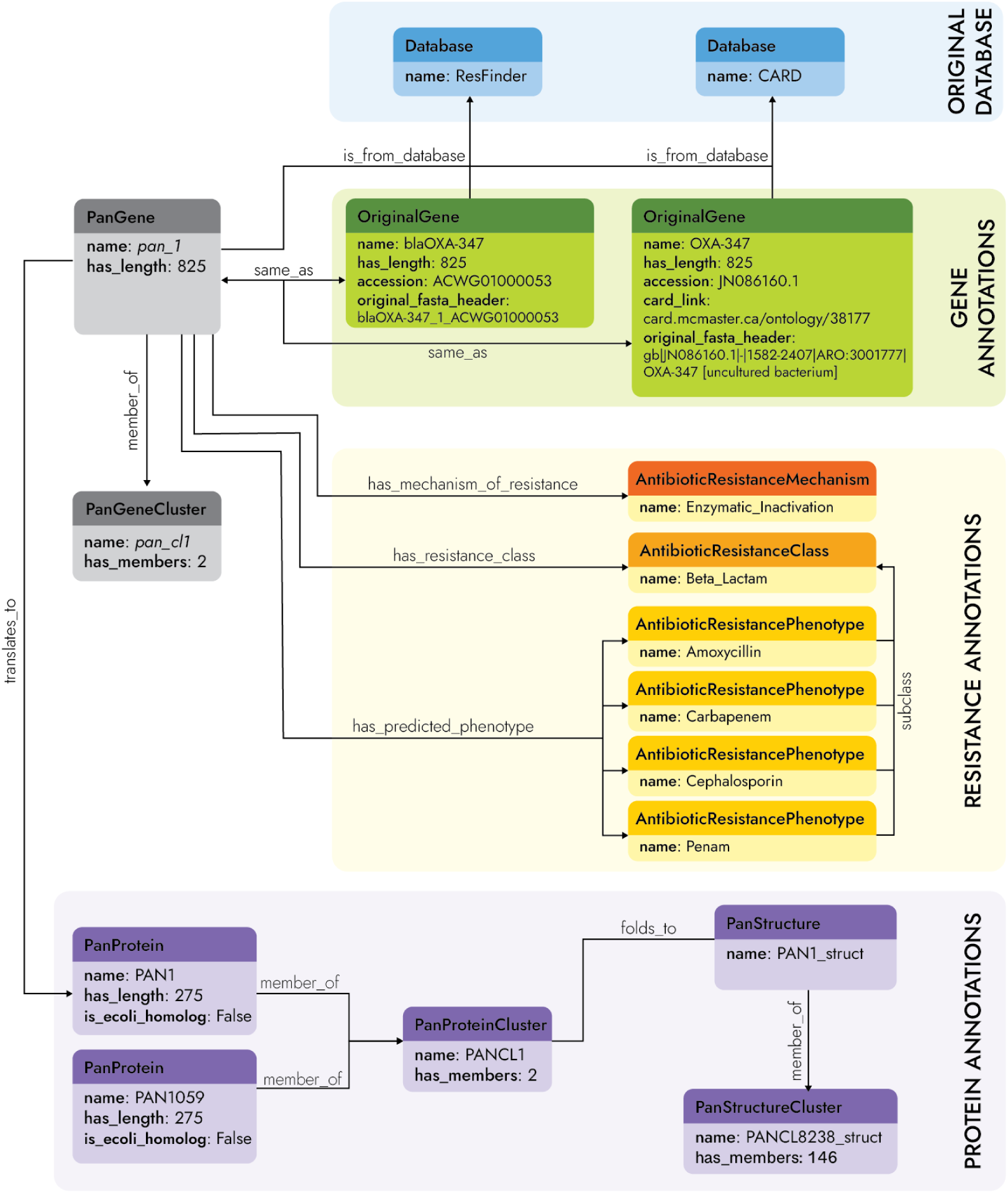
Schematic representation of the PanRes ontology. A single gene (*pan_1*) illustrates how information is defined through Classes (boxes) and Object/Annotation properties (arrows). The ontology connects original gene entries to the PanGene individual and links it to the corresponding resistance annotations (mechanisms, classes, and phenotypes). With the release of PanRes, genes are now also translated into proteins and predicted folds, and the information related to these individuals is likewise accessible in the ontology.

To flag resistance genes that may instead represent *E. coli* housekeeping homologs, we compared the PanRes protein sequences to a soft-core genome dataset (proteins present in >95% of *E. coli* strains) and identified 239 homologs. These are annotated in the ontology with an *is_ecoli_homolog* property, enabling users to identify and exclude sequences that could bias read-based AMR detection.

### Structure-based HMMs improve detection of remote homologs

To extend detection beyond conventional sequence similarity, we generated HMM profiles from the high-confidence structurally coherent PanRes clusters and compared these 3D-based searches with BLASTp across seven sewage metagenomic co-assemblies, testing whether they could recover more distant homologs. We considered only high-confidence hits (e-value < 1e−5), ignored matches to *E. coli* core proteins, and counted each query once by its highest-scoring retained hit.

Across all seven samples, on average 54.2% of hits were shared between methods, while 35.3% were uniquely identified by HMMER. Complete disagreements between the methods were rare (<0.3%), showing that BLASTp and HMMER usually assigned proteins to the same family when both methods detected them (Supplementary figure 2).

When HMM-unique hits were forced to BLASTp-align without thresholds, their sequence identity was lower than for queries detected by both methods (Supplementary figure 3). Proteins below the Rost length-identity curve fall in the twilight zone of sequence similarity. HMM-unique proteins were strongly enriched in this region, while proteins detected by both HMMER and BLAST mostly fell above it. In the Copenhagen RA sample, for example, 63.1% of HMM-unique proteins fell below the curve (Figure 4A), compared with only 4.6% of proteins detected by both methods (Supplementary table 7).

**Figure 4.**
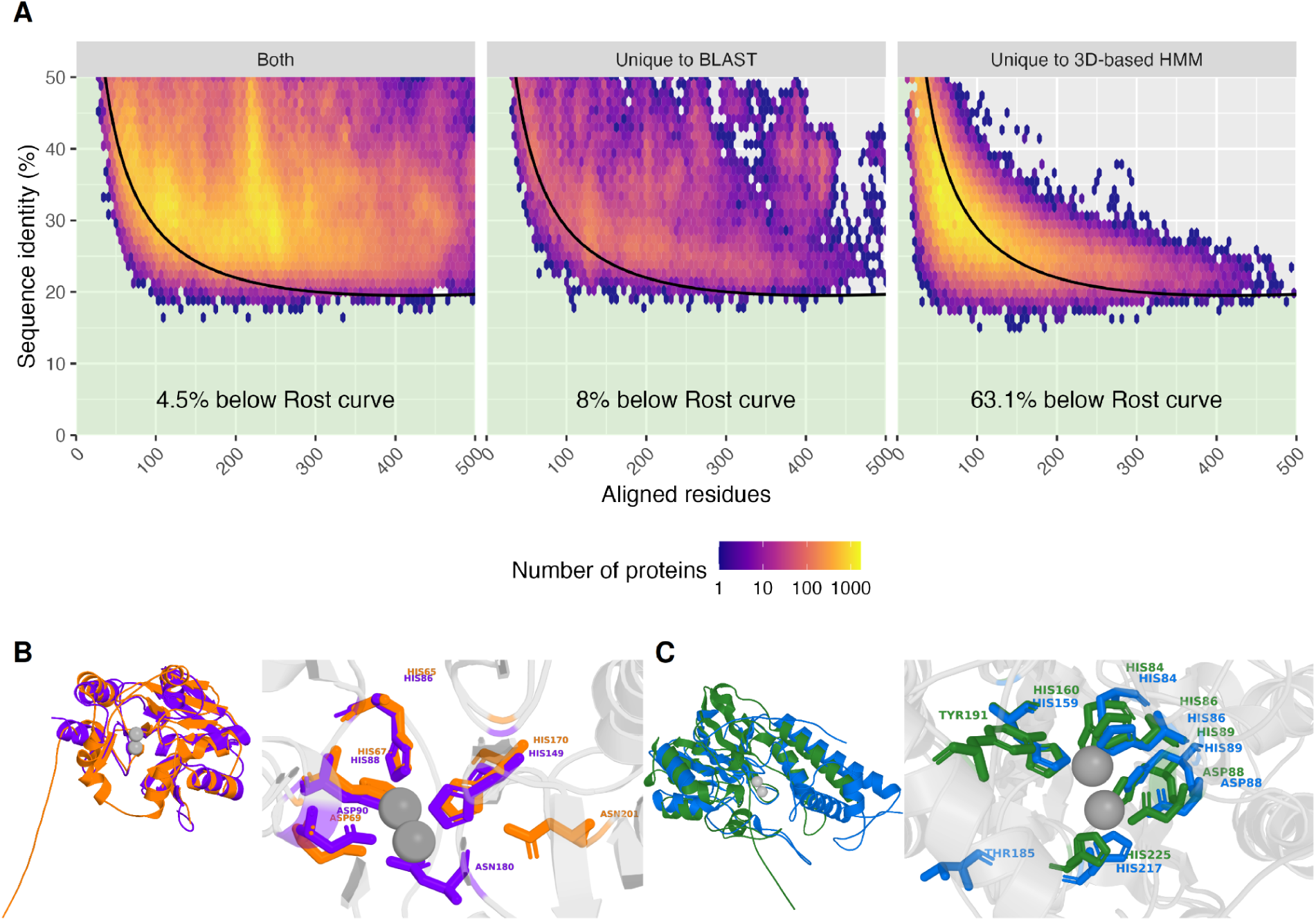
**A.** Sequence identity versus alignment length distribution of proteins detected by HMMER and BLASTp in Copenhagen RA. The area under the Rost protein-identity curve (green tint) contains proteins in the “Twilight zone” of protein homology^25^. The plot is cropped to 0–500 aligned residues and 0–50% identity to highlight low-identity hits. **B.** 3D alignment of hit protein *k127_37667511_19* (orange) to *PANCL12631_struct* reference structure of BcII (purple) obtained from the PDB database (1bc2). The active site residues around the Zn atoms (grey spheres) stay preserved. **C.** 3D alignment of a chemically conserved hit *k127_52446778_1* (blue) to the *PANCL5265_struct* reference structure (1sml) of metallo-beta-lactamase L1 type 3 (green). The active-site architecture is retained, including the Zn-coordinating residues, with a tyrosine-to-threonine substitution in place of a hydrogen donor.

To test whether HMM-unique proteins were entirely missed by BLAST or just fell below the high-confidence e-value threshold, we progressively relaxed the BLASTp threshold and examined whether these proteins could be recovered. Using the same top-hit filtering strategy as in the main comparison, some HMM-unique proteins were recovered only at more permissive thresholds, but on average over 41.9% remained unrecovered even at e-value ≤ 1. Allowing recovery by any non-homolog BLASTp hit reduced this fraction but did not remove it (Supplementary figure 4).

80.1% of HMM-unique proteins were also supported by stringent structure-informed validation, with confident Foldseek matches to PanRes structures requiring at least 80% coverage of the PanRes target structure across the datasets (Supplementary table 8).

Together, these analyses show that 3D-based HMMs expand the searchable homology space beyond BLAST and that many HMM-unique proteins have very weak or no detectable pairwise sequence similarity to PanRes entries. However, these additional hits should be interpreted as remote homologs or candidates, not confirmed resistance proteins.

### PanRes identifies functionally conserved putative beta-lactamases

Because the HMM search produced many candidate resistance proteins, we focused on beta-lactamases for detailed analysis. They formed some of the largest and best-described PanRes clusters, and their well-studied active sites let us test whether function was supported by conservation of the residues needed for beta-lactamase activity.

We annotated catalytic sites in four large beta-lactamase HMM clusters representing Ambler classes A-D, covering both serine beta-lactamases and metallo-beta-lactamases, which rely on different catalytic mechanisms^48^. For each HMM hit, we assessed whether the catalytic positions were exactly conserved or substituted by physicochemically similar residues. This reduced the broad HMM output to 2,066 proteins with active-site patterns consistent with beta-lactamase function, spanning identities as low as 20.1%, with 26.2% below 30% identity.

To illustrate the structural support behind these predictions, we selected representative candidates for closer inspection. The first example was a strict active-site match from the *PANCL12631_struct* HMM in the Budapest sample. This protein had a significant HMM match (e-value = 4e−09), retained all catalytic residues, and was not recovered by BLASTp even with relaxed criteria (e-value ≤ 1). We aligned the predicted structure to the reference BcII beta-lactamase structure used for active-site scoring (PAN6055). The catalytic residues remained positioned around the zinc ions, preserving the metallo-beta-lactamase active-site architecture (Figure 4B). AlphaFold also independently matched this candidate to a metallo-beta-lactamase superfamily protein from *Thermus thermophilus*, further supporting a beta-lactamase-like fold (Supplementary figure 5A).

The second example was a chemically conserved active-site match from *PANCL5265_struct* in the Copenhagen RL sample. This candidate retained the full catalytic site of the reference L1-type metallo-beta-lactamase (PAN7175), although one catalytic residue differed at the sequence level. The catalytic tyrosine was replaced by threonine, a conservative substitution that preserves the hydroxyl-containing side chain involved in hydrogen bonding and electrostatic stabilization^49,50^. Structural alignment showed that this residue occupied a similar position in the catalytic pocket, while the surrounding zinc-coordinating residues remained conserved (Figure 4C). AlphaFold provided further support for a beta-lactamase-like fold by matching the candidate to a beta-lactamase domain protein from *Burkholderia multivorans* (Supplementary figure 5B). BLASTp detected the protein only weakly (e-value = 0.004), compared with a much stronger 3D-based match (e-value = 4.9e−15).

Together, these examples show how structure-derived HMMs can identify highly divergent beta-lactamase-like proteins, while active-site conservation helps prioritize the candidates most likely to share a resistance function.

We next placed the 314 best-supported candidates from medium- and high-quality MAGs or plasmid contigs (Supplementary table 9) into the beta-lactamase phylogenies to compare them with known representatives and examine their taxonomic context. In the *PANCL12631_struct tree*, these hits were distributed across the B1/B2 metallo-beta-lactamase cluster and formed several hit-rich clades (Figure 5). Within the largest cluster, *Burkholderiaceae* was the most represented family among the retained hits, accounting for 50 candidates. This is notable because *Burkholderia*-related taxa are known for diverse secondary metabolism, including antimicrobial compounds, making them a plausible source of beta-lactamase-like diversity^51,52^. The plasmid-associated hits were distributed among chromosomal hits in the tree, showing that related beta-lactamase-like proteins may occur in both contexts.

**Figure 5.**
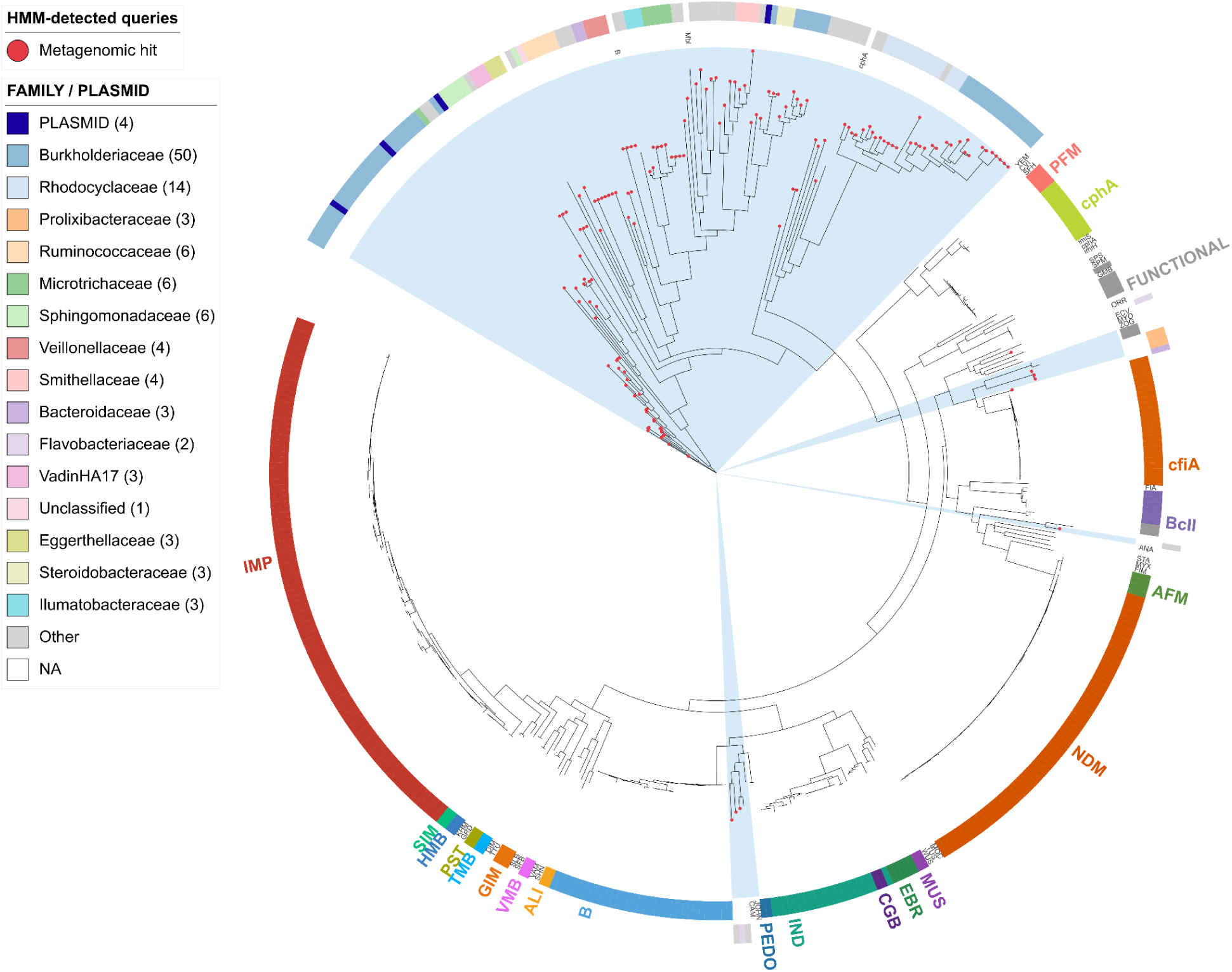
Maximum-likelihood tree of the PANCL12631_struct cluster, containing Ambler class B1/B2 metallo-beta-lactamases. The tree includes both PanRes reference proteins and HMM-detected sewage proteins from the PANCL12631_struct cluster. The inner ring shows beta-lactamase family annotations; FUNCTIONAL marks unnamed beta-lactamases from CsabaPal or ResFinderFG. The outer ring shows taxonomy at family level or PLASMID for plasmid-associated hits. Red circles indicate sewage hits, and blue shading marks hit-rich clades.

Several *PANCL12631_struct* hits were placed outside well-characterized clinical beta-lactamase groups and closer to latent or less characterized proteins. A similar phylogenetic spread was observed for the B3 metallo-beta-lactamase cluster (Supplementary figure 6), whereas the larger serine beta-lactamase clusters retained fewer hits after filtering (Supplementary figure 7–8). This likely reflects the difficulty of resolving highly diverse serine beta-lactamases from fragmented wastewater metagenomes. Because we only retained hits on long contigs in good quality bins, or on long plasmid-associated contigs, this conservative filtering may remove many partial or ambiguous candidates from highly diverse clusters. These results indicate that the retained candidates represent diverse beta-lactamase-like proteins with supporting active-site evidence, although experimental validation is needed to confirm resistance function.

## Discussion

PanRes was built to connect acquired and latent resistance determinants across antibiotic, biocide, and metal resistance with ontology-based annotation, predicted protein structures, and 3D-based HMMs.

A recent WHO-authored review identified several major challenges for AMR databases: resources are fragmented and overlapping, annotations are difficult to compare, and no single resource covers the entire AMR diversity^53^. PanRes addresses these challenges by integrating acquired AMR genes, biocide and metal resistance genes, and latent determinants identified through functional metagenomics into a single resource. Rather than replacing specialised databases, PanRes makes their content easier to use together by collapsing duplicate entries while keeping each gene traceable to its original name, accession, and source database. The shared ontology then links these across the resource. This makes it possible to ask questions that span the wider resistome rather than one database at a time.

The same review also stresses that AMR databases are limited by the diversity already known. Our results show this clearly for the latent resistome. Many functional-metagenomics genes had no close relatives among clinically recognised AMR genes, and latent-only clusters typically contained just a single sequence. This shows that much of the latent resistome remains poorly sampled, and that sequence-based searches are unlikely to capture its full diversity. Therefore, we designed the 3D-based HMMs to reach into this poorly sampled space.

Several AMR resources have already shown that HMMs can help improve detection of divergent resistance genes. BLAST-based searches work well for close homologs of known resistance genes, but lose sensitivity when proteins share only weak amino acid similarity with the available references. Resfams used curated protein-family HMMs to annotate translated metagenomic sequences and recovered resistance proteins missed by BLAST-based searches^54^. fARGene similarly uses optimized profile HMMs to identify and reconstruct divergent candidate ARGs directly from metagenomic reads^55^, achieving higher sensitivity for novel beta-lactamases than alternative approaches and experimental validation of 81% of tested predictions. ARGs-OAP v2.0 uses SARGfam HMMs to improve detection of remote homologs, while Meta-MARC applies DNA-based HMMs for AMR classification from reads or assemblies^56,57^. PanRes follows the same principle, but differs in how the profiles are built. The HMMs are not defined only by sequence families, AMR gene subtypes, or gene nomenclature. They are built from MSAs guided by structurally coherent protein clusters, anchoring the search in conserved protein folds rather than sequence similarity alone. This structural layer allowed PanRes to recover distant candidates enriched in the twilight zone of protein homology, including proteins not recovered by BLASTp even at relaxed thresholds.

Current AMR databases also differ in how cautiously they represent genotype-to-phenotype evidence. Because gene presence alone does not always imply resistance^58^, PanRes includes an *is_ecoli_homolog* annotation for 239 proteins, allowing users to exclude likely species markers from read-based analyses. We used *E. coli* as a well-characterized starting point, but this approach is limited to one species and will miss core proteins in other taxa. Expanding this comparison to a broader bacterial core-gene reference set will therefore be important in future versions of PanRes.

Finally, there are many usability barriers since many tools require bioinformatics expertise. The PanRes web interface lets users browse, filter, visualize and download the ontology, sequences and structures without command-line work, making the resource accessible to non-specialists.

While broader sensitivity is the main advantage of the PanRes 3D-based HMMs, it also means that the results need careful interpretation. A structural HMM hit supports remote homology, but it does not confirm resistance function. In our beta-lactamase analysis, we tried to reduce the uncertainty by adding active-site conservation as a filter. This made the candidate set more conservative, but probably also removed real beta-lactamases. Active sites can vary across large families, and metagenomic assemblies often fragment genes or place them on short contigs that failed our bin or plasmid filters. However, it is not possible to apply this strategy database-wide, as many mechanisms, such as efflux, target protection, or metal resistance, cannot be filtered by a small set of catalytic residues. Structure-based approaches such as Folddisco^59^ could provide an additional way to prioritize candidates when conserved functional sites are known.

A related structure-informed workflow was recently applied to beta-lactamases from Antarctic and Subantarctic soil metagenomes^17^, where permissive sequence clustering was followed by strict structural clustering to distinguish beta-lactamases from related but functionally distinct proteins. The retained candidates generally remained above ∼40% amino-acid identity to known references, consistent with the study’s conservative focus on high-confidence classification within beta-lactamases. Structure was therefore used as a late-stage filter after permissive sequence clustering. In contrast, PanRes uses structural similarity upstream to guide HMM construction across the full resistance gene collection, enabling detection of substantially more divergent proteins, including beta-lactamase candidates with sequence identities as low as ∼20% and retained catalytic residues.

The profiles also differ in how much information they can capture. Many structural clusters are singletons, and HMMs built from a single protein cannot learn family-level conservation in the same way as HMMs built from diverse alignments. This makes it harder to separate true remote homologs from false positives, especially in the latent resistome where many genes are still sparsely sampled. In this case, HMM searches are therefore unlikely to provide the same advantage over pairwise sequence comparison. Expanding these clusters as new homologs are identified will therefore be crucial. As the clusters grow, the HMMs will become more robust and specific, while also revealing more of the AMR diversity that is currently represented by isolated genes.

This is also why the PanRes models are best viewed as a discovery resource rather than a direct prediction tool for epidemiology or clinical decision making. Their broader sensitivity is useful for finding distant candidates, connecting acquired and latent resistance genes, exploring resistance-family diversity, and understanding where resistance came from. However, broad HMM hits should still be treated as candidates and require stricter filtering or experimental support before being used for surveillance.

Future versions of PanRes should focus on making the evidence behind each gene easier to interpret. One important step would be to separate tested negative phenotypes from missing phenotype information. At the moment, many databases do not clearly distinguish between a gene that was tested against an antibiotic and showed no resistance, and a gene that was simply never tested against that antibiotic. This matters because functional studies often test only a selected set of compounds, while the same gene may affect related antibiotics, biocides, or metals that were not included in the experiment. Some untested associations could be inferred from related compounds, structural clusters, active-site similarity, or closeness to characterized genes. Including observed positive, observed negative, not tested, and inferred phenotypes in future versions of PanRes would make the evidence behind each annotation clearer and help guide future research.

In conclusion, PanRes provides a structured resource for exploring AMR across database boundaries and supports discovery beyond close sequence matches. We hope it will serve as a useful resource for AMR researchers studying both known resistance genes and the wider, less characterized resistome.

## Data availability

The PanRes database and its related files have been added as an update to the original gene collection bundled with ARGprofiler (10.5281/zenodo.22876216). It contains the genes, 11,717 unique protein sequences, 1,703 protein folds, the newly added ontology and the HMM profiles. The code to generate the ontology is available at: https://github.com/rambiolab/PanResOntology/ and 10.5281/zenodo.23034831. The PanRes web interface is available at: https://panres.rambio.dk/. The sewage metagenomic reads, assemblies, and MAGs analysed in this study were generated in the original wastewater metagenomics study by Becsei et al. and are available on the European Nucleotide Archive under accession code PRJEB68319. The *E. coli* soft-core protein data used for homolog identification were obtained from the pangenome study by Tantoso et al. and are available at https://github.com/biierwint/ecoli_pangenome.

