## Supplementary information for "PanRes: A database of latent and acquired antimicrobial resistance allowing 3D-based protein homology search"

### Supplementary methods

#### Protein structure predictions

First, multiple sequence alignments were generated with colabfold_search using environmental sequences and structural templates (*--use-env 1 --use-templates 1*). The environmental, primary sequence, and template databases were *colabfold_envdb_202108*, *uniref30_2302_db*, and *pdb100_230517*, respectively. Structure prediction was performed with colabfold_batch using the *--use-gpu-relax* option, while all other settings were left at default: five predicted models per sequence, three recycles, 2,000 relaxation iterations, and no predicted Local Distance Difference Test (pLDDT) filtering. Structure prediction was performed in batches on NVIDIA Tesla V100 16GB GPUs, and the highest-ranking predicted structure for each protein was retained for downstream analyses.

#### Structure-based clustering and pairwise structural comparison

Foldseek1 was run in global alignment mode with TMalign (*--alignment-type 1*), and clustering was based on the alignment Template Modeling (TM) score (*--tmscore-threshold-mode 0*). Pairwise structural alignments were required to meet a minimum alignment coverage of both the query and the target structures (*--cov-mode 0*). All-vs-all pairwise comparison was performed using Foldseek's easy-search with the exhaustive search enabled (*--exhaustive-search*). Global TMalign alignment was chosen mirroring the clustering settings. No e-value cutoff was selected and other parameters were kept default.

#### Pfam cluster purity analysis

To assess the coherence of the 3D clusters, we screened all member protein sequences in each cluster against Pfam-A models[2] (v38.1) using HMMER[3] (v3.4) with default parameters. For each protein, we retained only the top-ranking hit, defined as the hit with the lowest e-value, resulting in one Pfam annotation per protein. Cluster-level Pfam consistency was then calculated as the proportion of annotated proteins assigned to the dominant Pfam family in that cluster. This resulted in a consistency measure (0-1) for each cluster, where 1 indicates complete Pfam agreement among all the cluster members.

#### Identification and annotation of core *E. coli* homologs in PanRes

PanRes contains resistance genes identified by functional cloning, namely from ResFinderFG and the CsabaPal gene collection. ResFinderFG is a collection of genes from 50 functional metagenomics studies, the majority of which were performed with *E. coli* as the model organism. Genes identified this way may reflect mutations or expression effects in native *E. coli* genes, rather than genuine transferable AMR determinants. Such genes are not suitable for homology-based resistance predictions for example in metagenomics and can be mere species indicators. They can however be important for AMR researchers and therefore remain annotated in the ontology.

#### Structure-informed assessment of HMM-unique queries with Foldseek

All protein queries detected only by HMM in the seven wastewater datasets were converted to predicted 3Di representations using ModernProst[4] (commit dd30f4d0), a lightweight protein language model that accelerates ProstT5[5]-based translation from amino acid sequence to Foldseek's 3Di structural alphabet. The predicted 3Di representations were searched against the retained high-confidence PanRes structures using Foldseek[1] *easy-search*. Matches with an E-value ≤ 1e-5 were considered confident. We first required at least 80% coverage of both the query and target (*-c 0.8 --cov-mode 0*). Because HMM-unique proteins may contain additional domains or sequence extensions, we repeated the search requiring 80% coverage of the PanRes target only (*-c 0.8 --cov-mode 1*). For each query, the hit with the lowest E-value was retained. The structural cluster of the best Foldseek hit was then compared with the PanRes structural cluster from which the detecting HMM had been built.

#### Beta-lactamase analysis: phylogeny trees

For phylogenetic analysis, protein sequences were grouped by HMM cluster, and aligned against the corresponding HMM profiles to preserve the structural information. The resulting MSAs were used to infer phylogenetic trees with IQ-TREE[6] (v3.1.1) using ModelFinderPlus[7] to select the best fitting model, 1,000 ultrafast bootstrap replicates, and 1,000 SH-aLRT replicates. The models used were Q.PFAM+I+R7, Q.PFAM+R6, Q.PFAM+R10, and JTT+R6 for *PANCL12631*, *PANCL5265*, *PANCL8238*, and *PANCL9032*, respectively.

The phylogenies were visualized and annotated in iTOL[8]. Beta-lactamase families were shown as colored arcs in the inner ring and taxonomic assignments as outer ring of the tree. Taxonomic assignments were not considered reliable for contigs classified as plasmids, which were labelled accordingly. Arcs were added when members of the same family formed a continuous group of adjacent tips in the circular tree layout. The minimum group size was adjusted for each tree according to tree size and density (k = 2–5). Metagenomic hits were marked with red symbols at the corresponding leaf tips. Clades in which metagenomic hits made up at least 60% of the leaves were highlighted.

### Description of supplementary tables

**Supplementary table 1:** Effect of Foldseek clustering thresholds on structural cluster composition and structural coherence. For each threshold combination, the table reports the number of clusters, cluster size distribution, singleton frequency, cluster-level structural quality scores, and pairwise intra-cluster similarity metrics. The columns are described below.

| Column | Description |
|---|---|
| coverage | Minimum required alignment coverage used for Foldseek structural clustering. |
| tmaln_threshold | Minimum alignment TM-score required for structures to be clustered together. |
| low_conf_filter | Indicates whether low confidence structures (mean pLDDT <70) have been filtered out. |
| total_clusters | Total number of structural clusters produced under the given coverage and TM-score settings. |
| mean_cluster_size | Average number of protein representatives per structural cluster. |
| max_cluster_size | Size of the largest structural cluster. |
| n_singletons | Number of structural clusters containing only one structure representative. |
| pct_singletons | Percentage of all structural clusters that are singletons. |
| mean_alnTM_cluster | Mean cluster-level alignment TM-score, calculated as the average structural similarity within each cluster and then averaged across clusters. |
| mean_LDDT_cluster | Mean cluster-level LDDT score, describing local structural agreement within clusters. Higher values indicate more locally similar structures. |
| min_mean_alnTM_cluster | Lowest mean alignment TM-score observed for any structural cluster. This indicates the least structurally coherent cluster under the given settings. |

| | |
|---|---|
| mean_alnTM_pairwise | Mean pairwise alignment TM-score across all within-cluster structure comparisons. |
| min_alnTM_pairwise | Lowest pairwise alignment TM-score observed between any two structures assigned to the same cluster. |
| min_cov_pairwise | Lowest pairwise alignment coverage observed between any two structures assigned to the same cluster. |

**Supplementary table 2:** HMM assignment performance across clustering and alignment settings. Summary of HMM benchmarking results for different structural cluster threshold combinations, with and without MAFFT --keeplength. For each setting, the table reports false negatives, false positives, and the proportion of proteins assigned to the correct structural cluster within the top one or top two HMM hits. The columns are described below.

| Column | Description |
|---|---|
| mafft_keeplength_enabled | Whether MAFFT was run with --keeplength when adding sequences back to the structure-guided alignment. |
| coverage | Minimum structural alignment coverage used when defining the PanStructureClusters used to build the HMMs. |
| tmaln_threshold | Minimum alignment TM-score used for Foldseek structural clustering. Higher values require stronger structural similarity between proteins in the same cluster. |
| false_negatives | Number of PanRes proteins that did not get assigned to their correct structural-cluster HMM. |
| false_positives | Number of PanRes proteins assigned to an incorrect structural-cluster HMM. |
| pct_within_top1 | Percentage of proteins for which the correct HMM was the top-scoring hit. |
| pct_within_top2 | Percentage of proteins for which the correct HMM appeared among the top two hits. |

**Supplementary table 3:** Major sequence, cluster and resistance annotation classes in the PanRes ontology and their descriptions.

**Supplementary table 4:** Object properties used to connect PanRes ontology individuals, including relationships between genes, proteins, structures, clusters, and resistance-related entities.

**Supplementary table 5:** Annotation properties used to describe PanRes ontology individuals with database-derived, genetic, and phenotypic metadata.

**Supplementary table 6:** Coverage of major PanRes ontology relationships and resistance annotations.

| Column | Description |
|---|---|
| entity | Ontology entity/class |
| relationship | Object or annotation property of an ontology entity/class |
| n_annotated | Number of entries of the given entity that carry the specified relationship or annotation. |
| n_total | Total number of relevant entries that can possibly carry that relationship. For resistance annotations, this is restricted to the corresponding antimicrobial, biocide, or metal resistance gene class. |
| pct_annotated | Percentage of relevant entries carrying the specified relationship or annotation. |

**Supplementary table 7:** Twilight-zone classification of detected queries. For each co-assembly, proteins were grouped as detected by both BLASTp and HMMER, unique to BLASTp, or unique to HMMER. The table reports the number of proteins in each group whose best BLASTp alignment fell below or above the Rost twilight-zone curve, together with the corresponding percentages. The columns are described below.

| Column | Description |
|---|---|
| query_group | Detection group assigned to each query protein, for example HMM-only, BLAST-only, or detected by both methods. |
| city | Sewage co-assembly or sampling location where the query proteins were detected. |
| n_total | Total number of query proteins in that detection group and city. |
| below_curve | Number of query proteins whose best BLASTp alignment fell below the Rost curve, placing them in the sequence twilight zone. |
| above_curve | Number of query proteins whose best BLASTp alignment fell above the Rost curve. |
| pct_below | Percentage of query proteins in that group and city that fell below the Rost curve. |

| | |
|---|---|
| pct_above | Percentage of query proteins in that group and city that fell above the Rost curve. |

**Supplementary table 8: Foldseek validation of HMM-unique queries.** For each wastewater co-assembly, the table shows how many HMM-unique proteins had Foldseek support under the two coverage settings, and how often the best Foldseek hit matched the same PanRes structural cluster as the HMM assignment.

| **Column** | **Description** |
|---|---|
| city | Sewage co-assembly in which the HMM-unique query proteins were detected. |
| coverage_criterion | Foldseek coverage criterion used for the search. |
| n_total | Total number of HMM-unique proteins in the co-assembly. |
| n_any_hit | Number of HMM-unique proteins with any Foldseek hit passing the coverage criterion. |
| pct_any_hit | Percentage of HMM-unique proteins with any Foldseek hit. |
| n_confident | Number of HMM-unique proteins with a confident Foldseek hit (E-value ≤ 1e-5). |
| pct_confident | Percentage of HMM-unique proteins with a confident Foldseek hit. |
| n_same_cluster | Number of proteins whose best confident Foldseek hit belonged to the same PanRes structural cluster as the detecting HMM. |
| pct_same_among_confident | Percentage of confident Foldseek hits assigned to the same structural cluster as the detecting HMM. |
| pct_same_of_all_hmm_only | Percentage of all HMM-unique proteins whose best confident Foldseek hit belonged to the same structural cluster as the detecting HMM. |

**Supplementary table 9: Beta-lactamase candidates retained for phylogenetic analysis.** HMM-detected beta-lactamase candidates with exact or chemically conserved catalytic-site matches that passed contig, MAG-quality, or plasmid-context filters. The table includes HMM assignments, catalytic-site match counts, contig information, genetic context, MAG quality metrics, and taxonomy. The columns are described below.

| **Column name** | **Description** |
|---|---|
| city | Sewage co-assembly where the candidate protein was detected. |

| | |
|---|---|
| hmm | PanRes HMM profile that detected the query protein. |
| query_id | Identifier of the predicted metagenomic protein detected by the HMM. |
| template | PanRes reference protein used for catalytic-site mapping. |
| catalytic_total | Total number of catalytic residues evaluated for the corresponding template. |
| catalytic_match_count | Number of catalytic residues exactly conserved in the query protein. |
| chemical_catalytic_match_count | Number of catalytic residues conserved either exactly or by chemically similar amino acid substitution. |
| contig_id | Identifier of the contig carrying the detected query protein. |
| length | Length of the contig carrying the detected query protein. |
| depth | Sequencing depth of the contig carrying the detected query protein. |
| genetic_context | Genomic context of the contig, classified as chromosome or plasmid. |
| bin | Identifier of the genome bin or MAG containing the contig. |
| completeness_best | Measure of the genome bin completeness based on the best model used. |
| completeness_general | Estimated completeness of the genome bin using the general completeness model. |
| completeness_specific | Estimated completeness of the genome bin using the specific completeness model. |
| completeness_model_used | Model used to estimate genome bin completeness. |
| contamination | Estimated percentage of contamination in the genome bin. |
| coding_density | Density of coding sequences within the genome bin, calculated as the ratio of coding bases to total bases. |
| contig_n50 | N50 value of contigs in the genome bin. |
| average_gene_length | Average length of genes in the genome bin. |

| | |
|---|---|
| genome_size | Total size of the genome bin in base pairs. |
| gc_content | Guanine-cytosine content of the genome bin. |
| total_coding_sequences | Total number of coding sequences in the genome bin. |
| mimag | Genome bin quality classification according to MIMAG standards. |
| kingdom | Taxonomic kingdom classification of the genome bin. |
| phylum | Taxonomic phylum classification of the genome bin. |
| class | Taxonomic class classification of the genome bin. |
| order | Taxonomic order classification of the genome bin. |
| family | Taxonomic family classification of the genome bin. |
| genus | Taxonomic genus classification of the genome bin. |
| species | Taxonomic species classification of the genome bin. |

### Supplementary figures

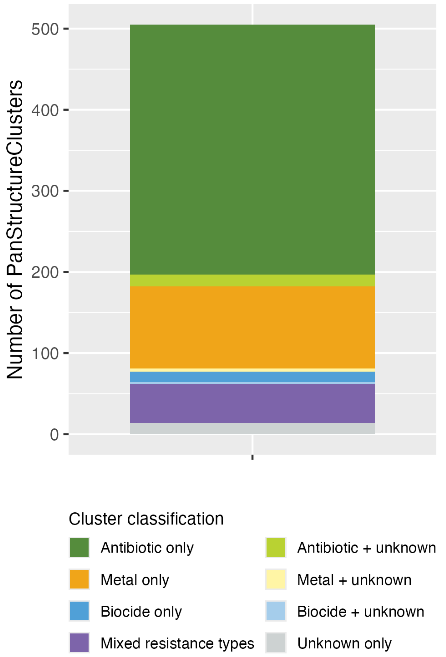


**Supplementary figure 1:** Antimicrobial resistance type composition of each PanStructureCluster (structural cluster).

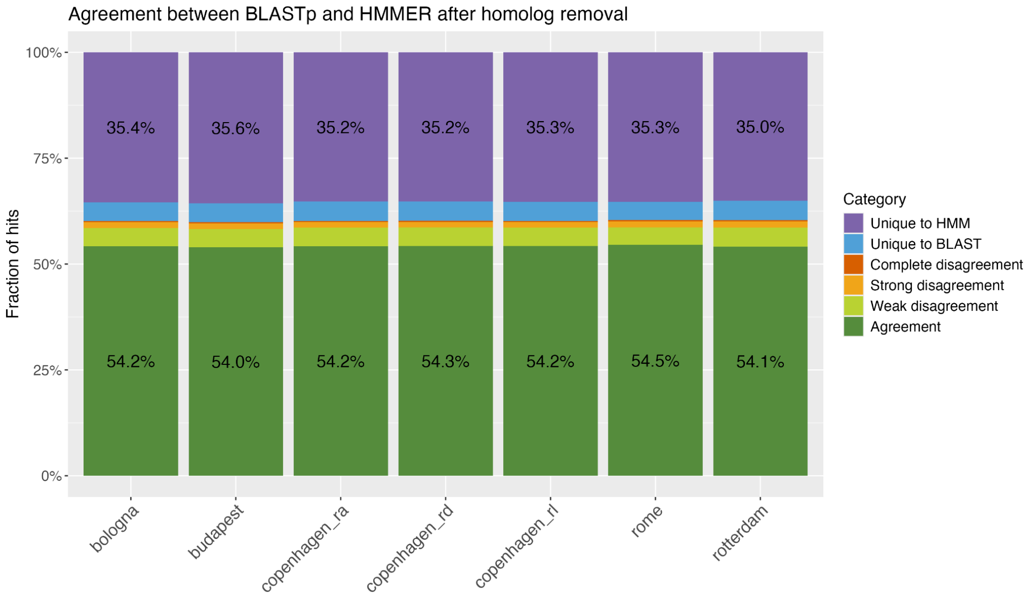


**Supplementary figure 2:** Agreement between BLASTp and HMMER query detection after *E. coli* core protein homolog removal. Stacked bar plot showing the fraction of query proteins assigned to each comparison category across the seven sewage co-assemblies. Agreement indicates that both methods selected the same target, while disagreement categories describe cases where the methods selected different top targets (see Methods).

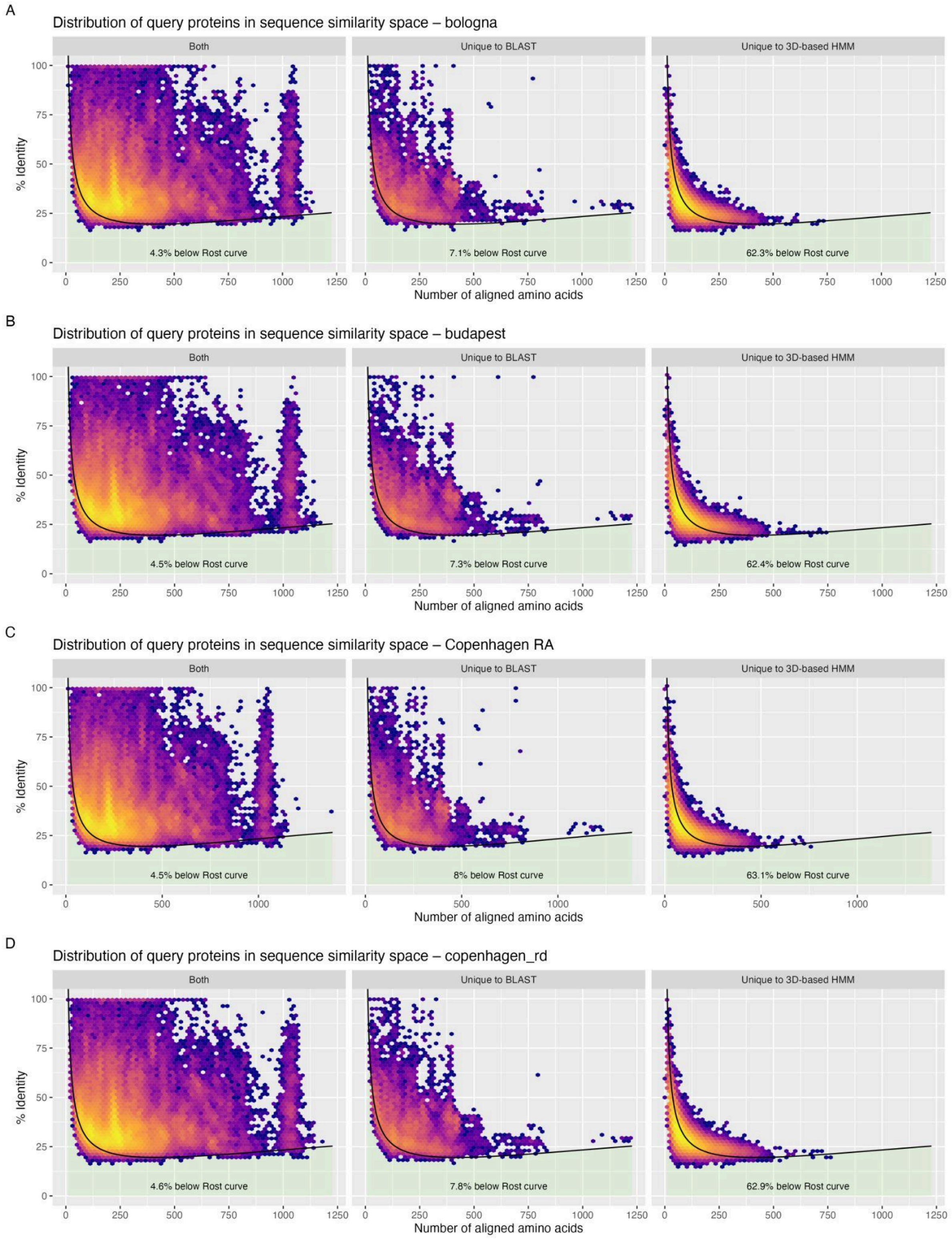
A
Distribution of query proteins in sequence similarity space – bologna
Both
Unique to BLAST
Unique to 3D-based HMM
% Identity
4.3% below Rost curve
7.1% below Rost curve
62.3% below Rost curve
Number of aligned amino acids
B
Distribution of query proteins in sequence similarity space – budapest
Both
Unique to BLAST
Unique to 3D-based HMM
% Identity
4.5% below Rost curve
7.3% below Rost curve
62.4% below Rost curve
Number of aligned amino acids
C
Distribution of query proteins in sequence similarity space – Copenhagen RA
Both
Unique to BLAST
Unique to 3D-based HMM
% Identity
4.5% below Rost curve
8% below Rost curve
63.1% below Rost curve
Number of aligned amino acids
D
Distribution of query proteins in sequence similarity space – copenhagen_rd
Both
Unique to BLAST
Unique to 3D-based HMM
% Identity
4.6% below Rost curve
7.8% below Rost curve
62.9% below Rost curve
Number of aligned amino acids

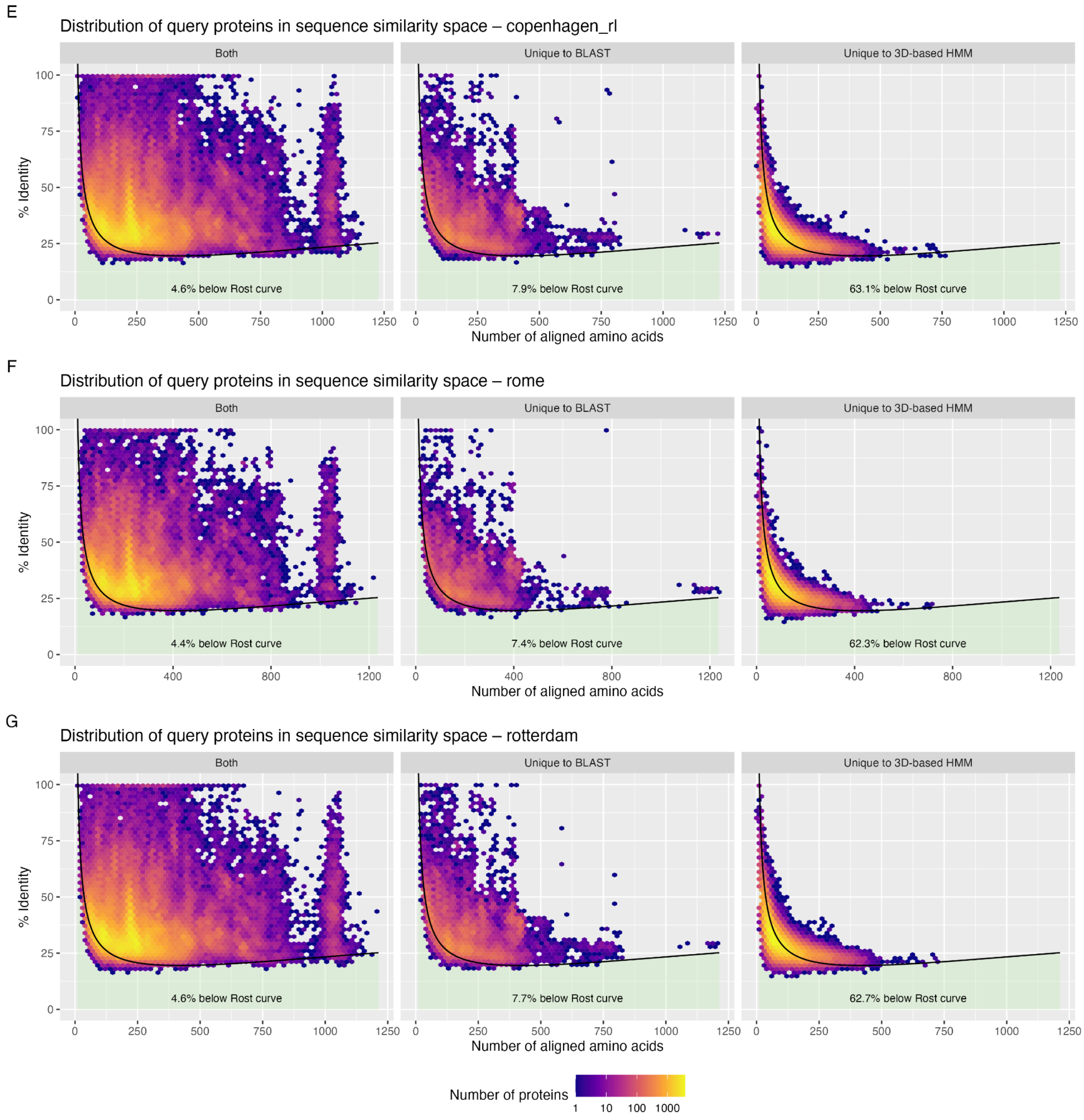


**Supplementary figure 3: Sequence identity versus alignment length distribution of proteins detected by HMMER and BLASTp in the seven co-assemblies.** Green shaded area indicates proteins in the "Twilight zone" of protein homology - area below the Rost protein identity curve. One best hit per query is shown.

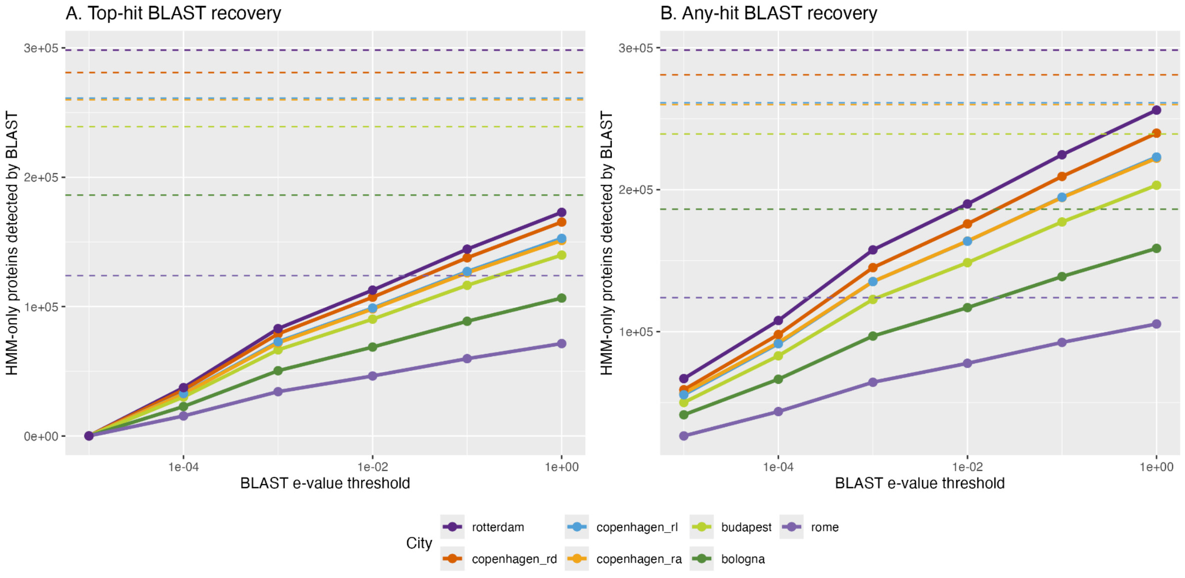


**Supplementary figure 4: Number of HMM-unique proteins recovered by BLASTp across increasingly relaxed e-value thresholds.** Recovery was evaluated using either **A.** the top BLASTp hit only or **B.** any non-homolog BLASTp hit. Dashed lines indicate the total number of HMM-unique proteins per city. On average, 41.9% remained unrecovered in A and 14.7% in B, at e-value ≤ 1.

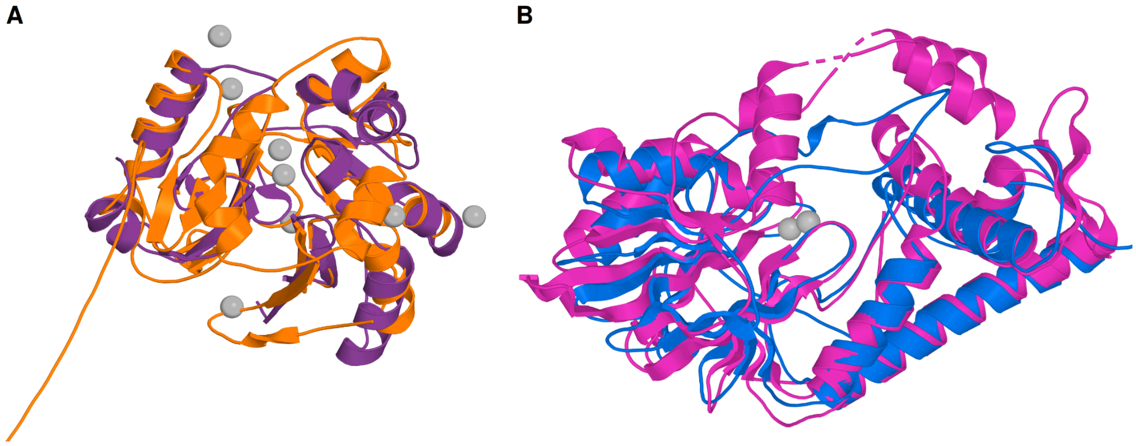


**Supplementary figure 5: Structural alignments supporting selected metallo-beta-lactamase candidates. A.** Predicted structure of the Budapest PANCL12631_struct strict active-site match *k127_37667511_19* (orange) aligned to a metallo-beta-lactamase superfamily protein from *Thermus thermophilus* (purple, PDB entry 2zwr). **B.** Predicted structure of the Copenhagen RL PANCL5265_struct chemically conserved active-site match *k127_52446778_1* (blue) aligned to a beta-lactamase domain protein from *Burkholderia multivorans* (pink, PDB entry 5u8o).

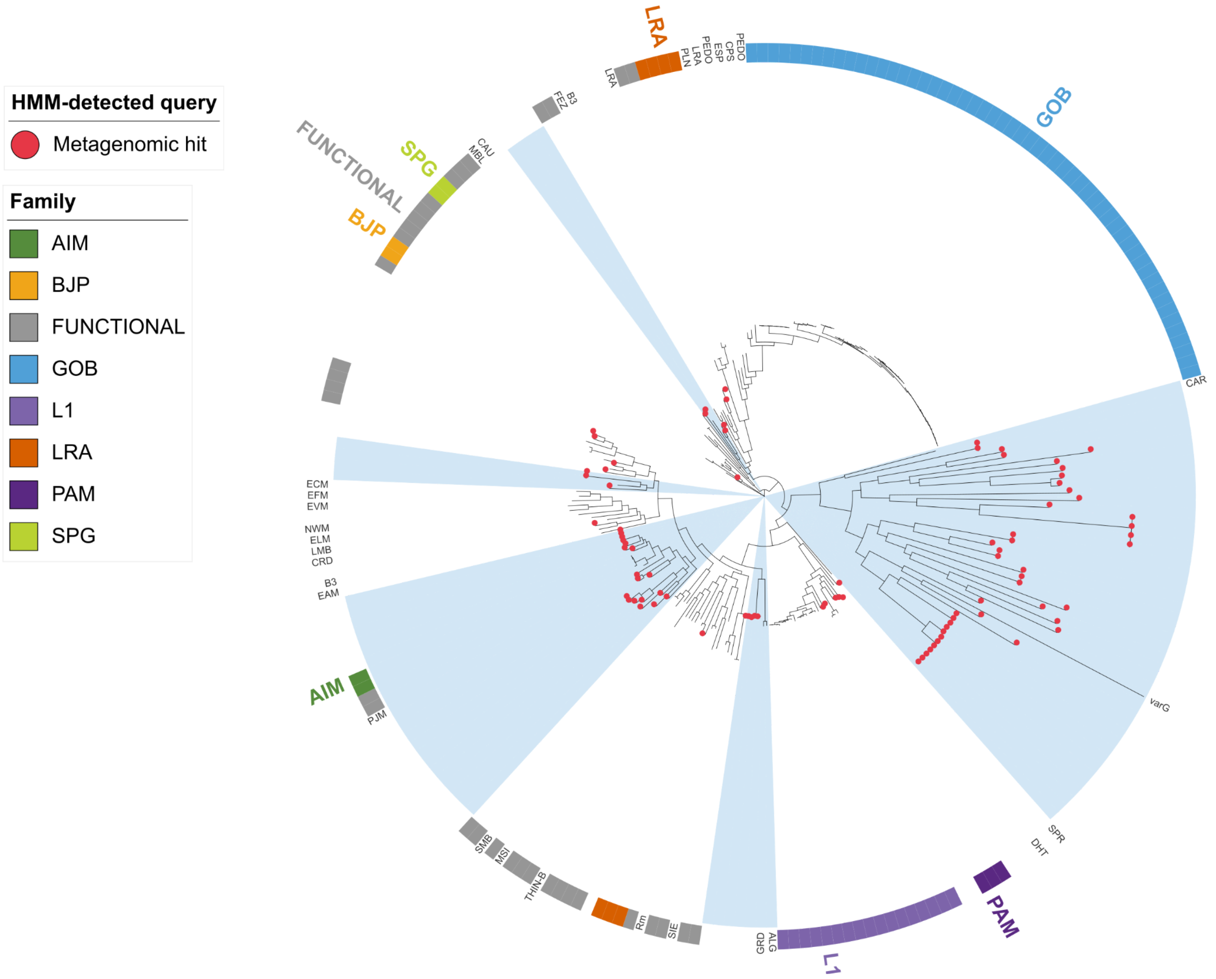


**Supplementary figure 6: Maximum-likelihood tree of the PANCL5265_struct cluster, containing Ambler class B3 metallo-β-lactamases.** The ring shows beta-lactamase family annotations, with unnamed beta-lactamases from CsabaPal/ResFinderFG marked as FUNCTIONAL. Red circles mark HMM-detected metagenomic hits from sewage co-assemblies.

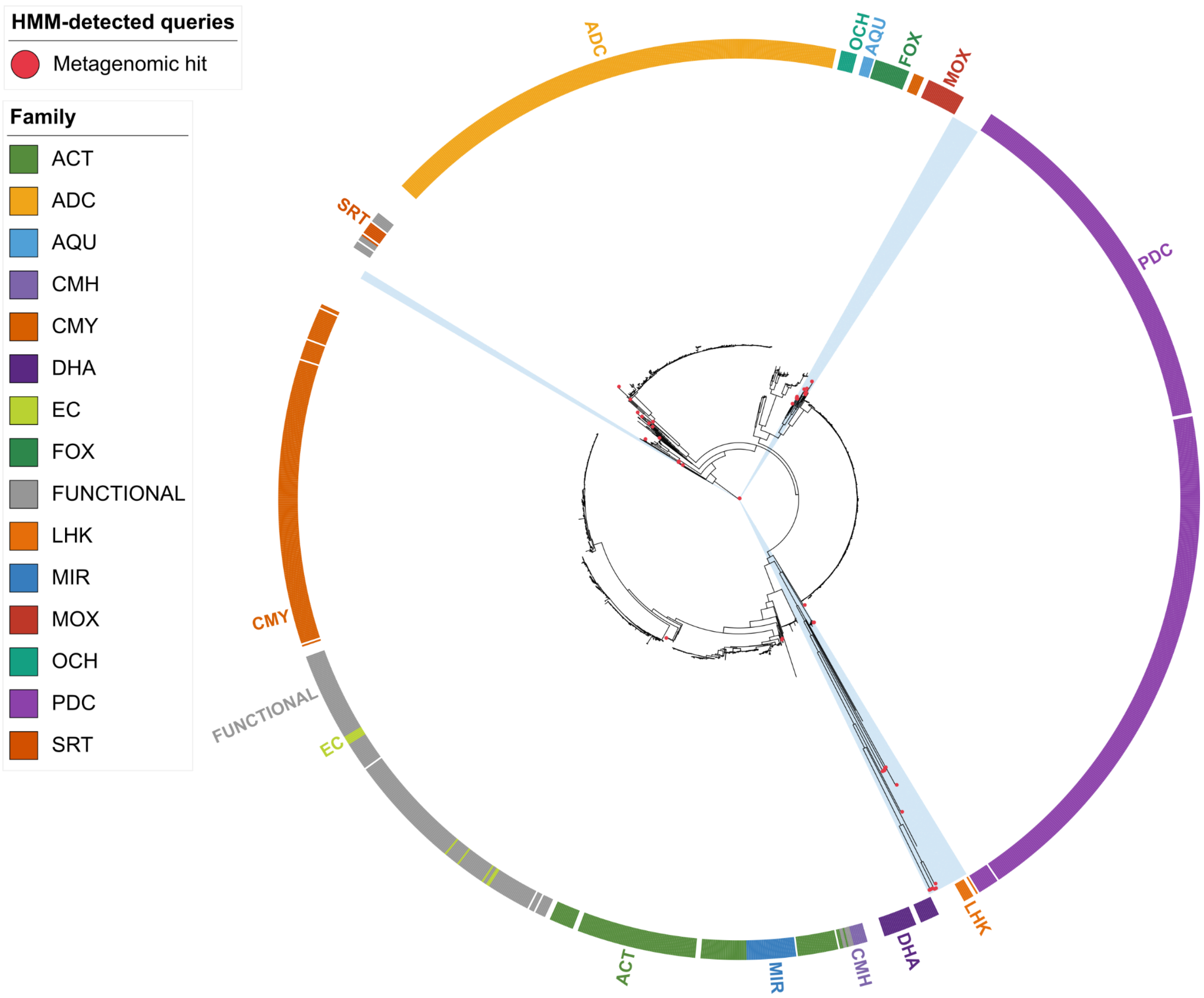


**Supplementary figure 7: Maximum-likelihood tree of the PANCL9032_struct cluster, containing Ambler class A/C serine β-lactamases.** The ring shows beta-lactamase family annotations, with unnamed beta-lactamases from CsabaPal/ResFinderFG marked as FUNCTIONAL. Red circles mark HMM-detected metagenomic hits from sewage co-assemblies.

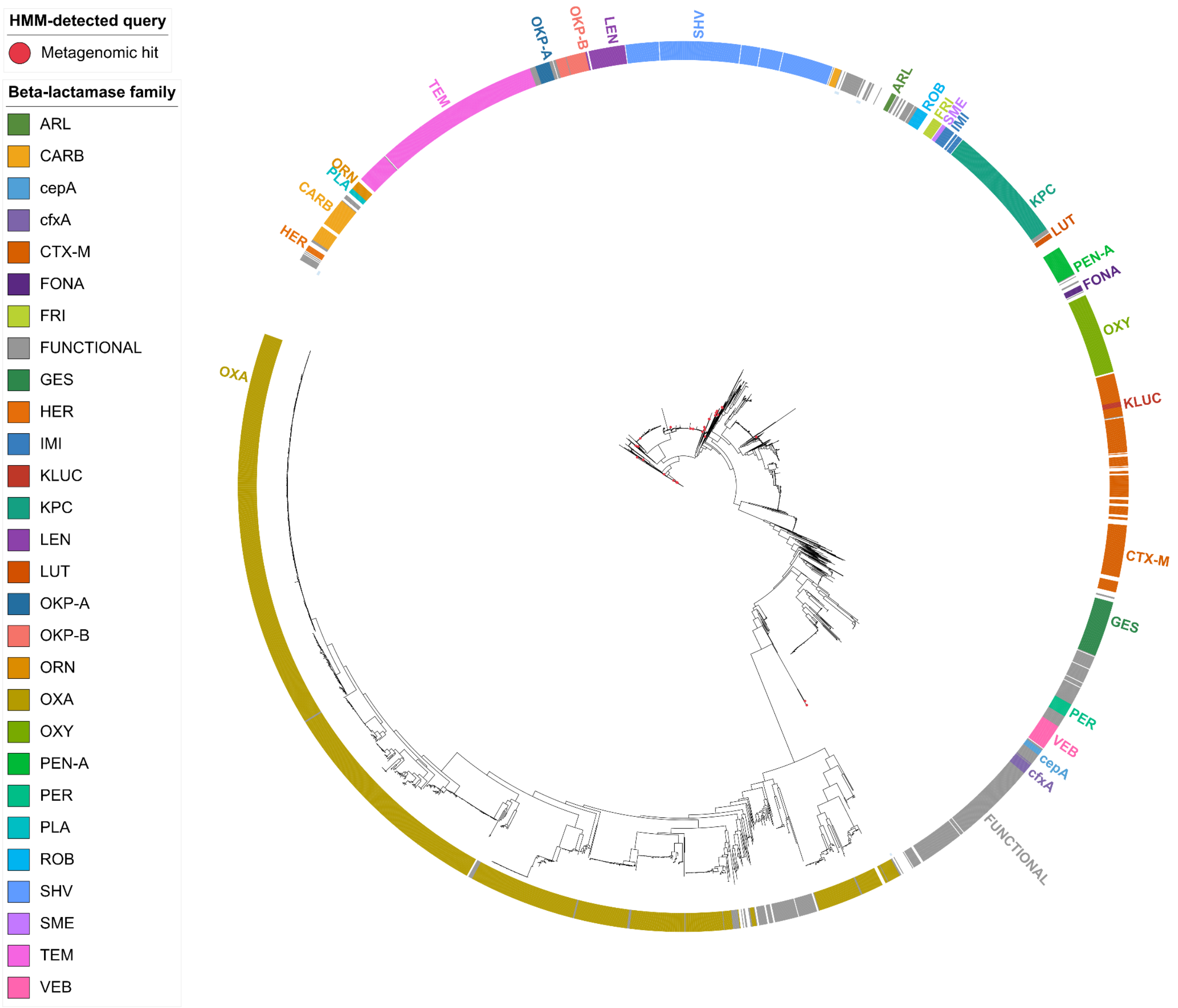


**Supplementary figure 8: Maximum-likelihood tree of the PANCL8238_struct cluster, containing Ambler class A/D serine β-lactamases.** The ring shows beta-lactamase family annotations, with unnamed beta-lactamases from CsabaPal/ResFinderFG marked as FUNCTIONAL. Red circles mark HMM-detected metagenomic hits from sewage co-assemblies.
